# COSMOS: A FAIR-aligned infrastructure for clinical trial data validation, warehousing, and interactive discovery

**DOI:** 10.64898/2026.07.16.738871

**Authors:** Charlie AK Roberts, Anna Nilsson-Takeuchi, Charlotte Stuart, Martin Chivers, Patricia Soares, Vincent Foure, Gareth Griffiths, Umar Niazi

**Author notes:** To whom correspondence should be addressed*. **Corresponding author’s email address:**. Authors contributed equally to this work (Charlie AK Roberts, Anna Nilsson-Takeuchi).

## Abstract

1.

**Summary:** The Clinical Omics System for Metadata and Outcome Storage (COSMOS) is an open-source, FAIR- and GCP aligned clinical trial unit (CTU) infrastructure designed to streamline the transition of academic clinical and multi-omic trial datasets into curated, analysis-ready repositories within Secure Data Environments (SDEs). By integrating an automated Data Quality and Data Validation (DQ&DV) “Trust Layer” with a relational Structured Query Language (SQL) schema, COSMOS enables programmatic and interactive data access via R Shiny applications. Dynamic integration of omics and clinical data is achieved through analytical data structures (e.g. ExpressionSet objects) linked via relational database identifiers. This lowers technical barriers for researchers and promotes governed data reuse and secondary discovery.

**Availability and Implementation:** COSMOS is freely available under the GPL-3.0 license at https://github.com/at2e19/SCTU_COSMOS_DQDV_Shiny

Supplementary information: Supplementary data are available at Bioinformatics online.

## 2. Introduction

Clinical trials generate two distinct but complementary data streams. Clinical data are systematically collected to address predefined primary and secondary outcome endpoints (e.g. survival and toxicity), whereas translational data (e.g. omics) are often generated after trial completion and primary outcome reporting. These data streams are rarely harmonized, resulting in disconnected “data lakes” that limit integration and downstream reuse, such as use by the discovery community that will result in hypothesis generation for future biomarker-guided clinical trials. COSMOS addresses this gap in the academic clinical trials unit setting by enabling structured linkage and reuse of both clinical and translational data within a unified framework, thereby extending the scientific value of trial datasets beyond their original scope.

As a Cancer Research UK core-funded Clinical Trials Unit (CTU), our vision at Southampton CTU (SCTU) is that discovery is at the heart of everything we do and so have a remit to have in-house, secure and responsible sharing of our trial data promoting discovery and open science in line with public mandate. To enable this, SCTU developed COSMOS (Clinical Omics System for Metadata and Outcome Storage) building upon previously introduced the SQL schema as a foundation for managing integrated datasets (Niazi *et al*. 2025). Such infrastructure depends entirely on the reliability of the underlying data. To address this, we developed the DQ&DV Trust Layer, informed by data quality and validation principles outlined by McGilvray (2021) to ensure an audit trail and data integrity prior to ingestion into the SQL database warehouse.

Furthermore, complex relational repositories traditionally exclude researchers who lack programming or SQL expertise. Although this database supports access programmatically e.g. through statistical software such as R, our suite of R shiny tools promote reproducible research and data democratisation by providing intuitive interfaces for “behind the glass” analysis within SDEs. Recognized by the Association for Clinical Data Management (ACDM) 2025 Innovation Prize (ACDM 2025), the COSMOS ecosystem (Figure 1) provides a Findable, Accessible, Interoperable, Reusable (FAIR)-aligned (Wilkinson *et al*. 2016), scalable, low-cost paradigm for clinical and translational data integration and governance that adheres to Good Clinical Practice (GCP) ICH E6(R3)(ICH Expert Working Group 2025) standards for the data life cycle.

**Figure 1:**
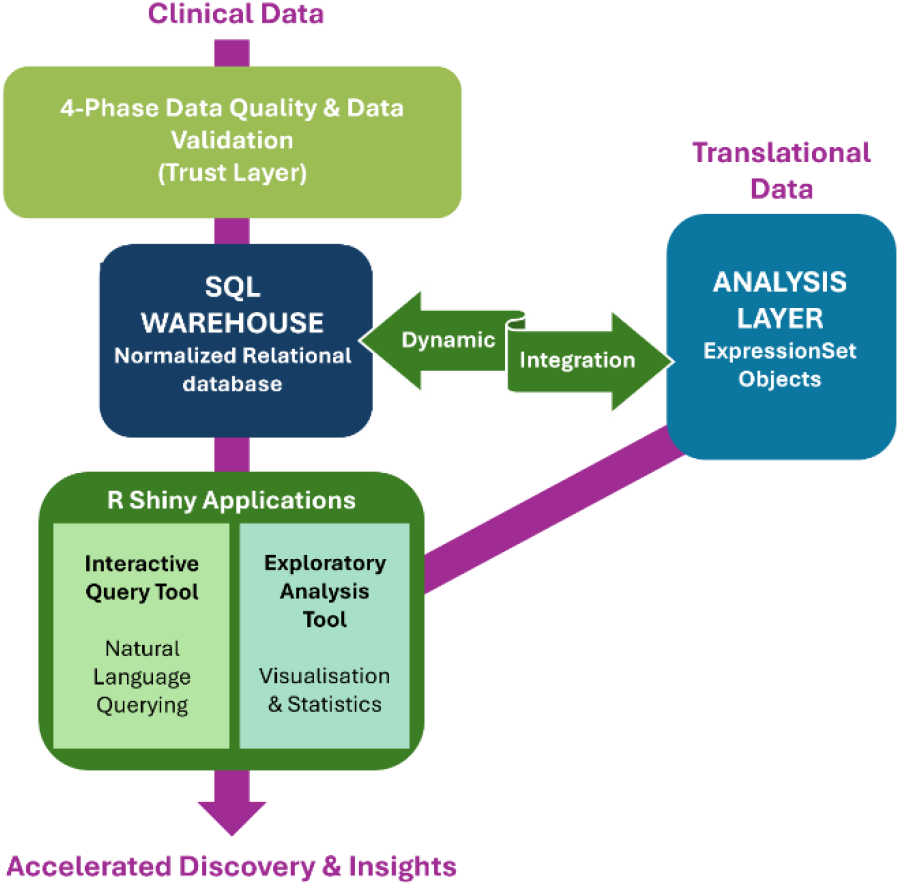
COSMOS architecture for clinical-omics data integration and discovery. Clinical data are processed through a four-phase Data Quality & Data Validation trust layer prior to ingestion into a normalized relational SQL warehouse. Translational datasets are processed using established and widely accepted analytical workflows. Analysis-ready datasets are constructed in an intermediate analytical layer as ExpressionSet objects, where omics measurements are dynamically linked to clinical variables through sample-level identifiers mapped to database keys. This design enables integrated, reproducible analyses without denormalising the underlying schema. Secure R shiny interfaces provide controlled, no-code access to querying and exploratory analysis within a secure data environment

## 3. Materials and Methods

### 3.1 The DQ&DV Pipeline: A Four-Phase Trust Layer

The DQ&DV pipeline implements an auditable migration from the Raw Data Lake (e.g. electronic data capture exports) to the SQL warehouse. This workflow operationalizes the Ten Steps to Quality Data framework (McGilvray 2021) through four distinct phases:

- **Phase 1: Data Discovery & Rule Derivation:** This phase consolidates core demographic and participant status data (e.g. eligibility, randomisation, and end-of-study data) to generate a definitive Master Subject List. A hierarchical status logic is applied to resolve inconsistencies across data sources. For example, where a participant is marked as both randomized and end-of-study due to conflicting records, predefined rules prioritise the most definitive trial milestone (e.g. end-of-study status overrides randomisation status). In parallel, a Data Dictionary is programmatically derived from trial metadata. This dictionary defines variable types, expected ranges, mandatory fields and Critical Data Elements (CDEs), forming the source of truth for downstream validation. CDEs are defined in consultation with key stakeholders, including trial managers, clinicians, and statisticians, ensuring alignment with downstream analyses such as statistical modelling and machine learning workflows.
- **Phase 2: Pre-Ingestion Quality Evaluation:** Raw datasets are first standardized through automated preprocessing, such as harmonisation of anonymised participant identifiers, trimming of whitespace and conversion of empty values to missing. Data are structured into participant-visit units of analysis to provide a consistent analytical framework. Each CDE undergoes systematic validation, including:
  - Structural QC: Completeness checks, identifying missing values in required fields.
  - Sematic QC: Data type validation, ensuring alignment with dictionary-defined types (e.g. numeric vs text mismatches).
  - Statistical QC:
    - Range and outlier detection based on empirical distributions of observed values.
    - Categorical consistency checks, flagging unexpected or infrequent category labels.
  Validation is programmatically performed across all relevant data, with each detected issue recorded alongside participant identifier, visit, variable name and issue type. Results are compiled into structured reports with both detailed issue logs and aggregated summaries, enabling review and correction by data managers prior to database ingestion.
- **Phase 3: Database Ingestion:** Validated participant and clinical data are ingested into the database according to the SQL schema defined previously in Niazi *et al*. 2025. The ingestion process preserves relational integrity by organising data into a standardized long-format structure indexed by participant and visit. A check-before-write mechanism ensures that exiting records are not duplicated, supporting safe iterative updates as source data evolve.
- **Phase 4: Post-Ingestion QC Certification:** A final reconciliation step ensures concordance between source data and warehouse. A random subset of data points is manually verified by data managers against source records. Successful verification results in formal certification of the dataset for downstream discovery use.

### 3.2 Linking Clinical and Translational Data

Following clinical data validation and warehouse ingestion, translational data modalities, such as transcriptomics, proteomics, and metabolomics, are subjected to quality control and processed using established, standardized analytical approaches appropriate for each modality prior to integration (e.g. Sumner *et al*. 2007, Conesa *et al*. 2016, Al-Amrani *et al*. 2021). Quality control procedures include assessment of batch effects, identification of unexpected sample clustering using dimensionality reduction techniques, and evaluation of normalization through inspection of summary statistics. The processed datasets are then incorporated into the SQL schema via a sample-linkage process. Metadata tables capture relationships between participants, visits, sample collection events, sample identifiers, assay platforms, and processed molecular matrices. Each translational sample is linked via foreign keys to a specific participant-visit unit of analysis, while individual sample-level metadata are preserved.

Analysis-ready datasets are constructed as Level 1 ExpressionSet objects (Gentleman *et al*. 2004), linking matrix-like omics data (e.g. gene expression count matrices with P features × N samples) to clinical variables and sample metadata through database linkage identifiers, enabling dynamic integration of clinical and omics data without denormalising the underlying database schema. This supports reproducible downstream analyses. ExpressionSet was selected for its simplicity and widespread compatibility with established Bioconductor workflows. The framework, however, remains compatible with more complex containers (e.g. MultiAssayExperiment, Ramos *et al*. 2017) as analytical requirements evolve.

### 3.3 Secure Discovery via R Shiny Framework

To democratize discovery while maintaining strict governance, we developed two R Shiny applications hosted in a secure SDE/Virtual Machine (VM) infrastructure. This enables authorized users to interrogate curated clinical and translational datasets without requiring direct database access or bespoke code.

1. **Interactive Query Tool:** This application allows users to retrieve data using predefined SQL queries. User-entered keywords (e.g. “available studies” or “clinical data”) are mapped to predefined queries via an inverted-index (Manning *et al*. 2008), enabling non-programmers to navigate the data and export CSV summary reports without requiring direct interaction with the underlying database.
2. **Exploratory Analysis Engine:** This module enables analysis of Level 1 omics data stored as *ExpressionSet* objects, with the ability to dynamically link relevant clinical data for integrated analysis. The tool provides exploratory, statistical and machine learning workflows, including summary statistic plots, hierarchical clustering, Principal Component Analysis, limma-based differential abundance analysis, and Random Forest-based feature ranking for discriminatory biomarker selection. This not only allows users to perform combined clinical-omics analyses within a single framework but does so from a no-code interface promoting data democratisation.

### 3.4 Security & Environment Awareness

Access to COSMOS is managed through a hybrid login pattern. The applications utilise system-defined Data Source Names (DSN) for secure authentication in production environments and manual credentials for local development. Furthermore, the framework uses environment-aware path configuration, ensuring seamless transitions between local workstation debugging and production VM environments. Access is restricted to authorized users, with all user access logged for audit purposes. Permissions are granted through controlled governance processes, such as formal data access requests to the Southampton CTU Data Sharing Committee or ticketing systems (for internal users). Raw underlying data are not directly accessible or exportable; instead, users can only export aggregated summaries or analysis outputs. Should a data sharing agreement allow for it, security controls can be configured so that data can be downloaded.

## 4. Results

### 4.1 Implementation in the CONFIRM Trial

The DQ&DV workflow was applied to the Cancer Research UK (CRUK)-funded CONFIRM trial that was conducted by SCTU (Fennell *et al*. 2021), comprising 366 UK participants investigating the impact of immunotherapy in mesothelioma with longitudinal data collected across 36 timepoints. A total of 5,557 participants– timepoint units of analysis were identified across all data. Phase 2 assessment identified no significant data quality issue, with all critical validation checks passing successfully. Automated screening highlighted a small number of participants with ages substantially lower than those observed in the wider cohort. Subsequent review confirmed these records to be valid, demonstrating the utility of the DQ&DV workflow in identifying unusual but valid observations for further review. The resulting integrated clinical and translational dataset is subsequently made available within a secure VM environment, enabling controlled access for both internal researchers and external collaborators.

### 4.2 Reproduction of Clinical Evidence

The utility of the Exploratory Analysis module was validated by reproducing results from a published metabolic signature study (McDonnell *et al*. 2025), conducted using data generated through the CRUK-funded DEPEND trial (Afolabi *et al*. 2022). The application linked plasma metabolomics with clinical metadata to perform differential abundance analysis and Random Forest for group-wise comparisons between pancreatic ductal adenocarcinoma (PDAC) patients and healthy volunteers. Pearson’s correlation analysis demonstrated a highly significant concordance (r= 0.987, p=7.85e^-15^) between the log fold changes (logFC) of the top differentially abundant metabolites identified through the R Shiny applications and bespoke analysis pipelines. By successfully replicating the metabolic signatures associated with resectable PDAC, this demonstrates that COSMOS provides expert-level analytical rigor within an accessible, no-code environment.

## 5. Discussion

Clinical trials increasingly generate clinical and molecular datasets that represent valuable resources for translational and discovery research beyond their original study objectives. The management and reuse of such data require robust governance, auditability and confidentiality safeguards. While repositories such as the European Genome Phenome Archive (Freeberg *et al*. 2022) provide secure storage and controlled access for sensitive omics and clinical trial datasets, they do not inherently support integration of molecular and clinical information for downstream exploratory analyses.

COSMOS addresses this gap by providing an open-source, low-cost, GCP-(ICH Expert Working Group, 2025) and FAIR-aligned framework (Wilkinson *et al*. 2016) for transforming heterogeneous trial data into validated, analysis-ready resources. The DQ&DV pipeline establishes an auditable trust layer before ingestion into a standardized SQL warehouse, while R Shiny applications enable authorized users to query and analyse linked clinical and molecular data without direct programming or database expertise. This enables secure, “behind-the-glass” discovery while preserving governance over participant-level data after the primary clinical outcome results have been published for trials.

Future work will extend COSMOS towards federated deployment, adopting shared schema and validation standards for multi-centre discovery while allowing institutions to retain local control of datasets.

## 6. Acknowledgments

