## Supplementary Materials for "COSMOS: A FAIR-aligned infrastructure for clinical trial data validation, warehousing, and interactive discovery"

**COSMOS summary**

A metadata-driven ingestion framework that converts source data (data lake) into a validated data warehouse through a combination of structural, completeness, semantic and statistical quality checks, data ingestion validation, and human review, complete with no-code analyses and query interfaces to explore the data.

**The DQ&DV Protocol: Systematic Trust Layer Implementation**

The COSMOS Data Quality and Data Validation (DQ&DV) pipeline is implemented as a series of modular R scripts. Each script generates a timestamped HTML report and a detailed text log, ensuring end-to-end traceability from raw ingestion to database release.

The DQ&DV pipeline is detailed below. A walkthrough of each script with a demo dataset can be found at the following GitHub Repository: https://github.com/at2e19/SCTU_COSMOS_DQDV_Shiny

**Phase 1: Data Discovery & Rule Derivation**

The first phase focuses on establishing a single, chronologically consistent record for every participant (Master Subject List), defining the official validation rules (Data Dictionary) and identifying Participant-Visit units of analysis.

*Master Subject List (Script 01)*

Consolidates core demographic and participant status data from source files (Primary Record, Eligibility, Randomisation, and End of Study) to form the master subject list. A hierarchical status algorithm resolves participant disposition assigning a single definitive status to each participant to establish the "Source of Truth" for participant counts. The hierarchical logic prioritises End of Study status (Completed/Withdrawn), followed by Not Eligible (Screen Failed), then Randomized, and defaulting to Consented if none of the preceding criteria are met. Automated data quality checks ensure the list meets critical standards for uniqueness, temporal consistency, and logical consistency. The final, validated Master Subject List is saved, providing a single source of truth for all downstream analyses.

*Data Dictionary Derivation (Script 02)*

Programmatically extracts metadata from the trial’s Architect Loader Specification to define expected data types, coded values, and longitudinal visit structures to form the Data Dictionary. It serves as the single source of truth for validation rules. A Critical column is added and manually populated to explicitly flag high-priority data (Critical Data Elements) elements for targeted quality control in the following phases of DQ&DV.

*Participant-Visit units of analysis (Script 03)*

Scans all clinical data to identify Participant-Visit units of analysis. Records lacking timepoint information are standardized by assigning a “cross-sectional” label, ensuring inclusion of non-visit-specific data. The output consists of a deduplicated list of Participant-Visit pairs for downstream processing, alongside a user-editable file requiring manual assignment of visit sequence according to the study protocol, enabling correct chronological ordering of study visits.

### Phase 2: Pre-Ingestion Quality Evaluation

This phase performs automated quality checks to identify potential major data quality issues prior to data ingestion. It leverages the Data Dictionary and the validated Master Subject List from Phase 1.

*Participant-Visit Overlap Heatmap (Script 04)*

*Did the expected visits occur?*

Generates a Participant-Visit incidence matrix and corresponding heatmap to visualize data completeness across study visits. Using the extracted Participant-Visit units of analysis, manually defined folder sequence, and Master Subject List, it constructs a binary matrix (presence/absence) with participants ordered by study status and visits arranged chronologically. The resulting heatmap provides an intuitive overview of visit level data coverage and highlights missing data patterns. Both the ordered matrix (CSV) and heatmap visualisation (PNG) are exported for further analysis and reporting.

*Participant-Visit Data Field Count (Script 05)*

*How much information exists in each visit?*

Quantifies data completeness by calculating the number of non-missing data for each participant at each study timepoint across all data. It aggregates these counts into a Participant-Visit matrix, visualized as a heatmap (Supplementary Figure 1) to represent data density, alongside summary bar plots of total data stratified by visit and participant. In parallel, the script enriches the data elements dictionary by adding these metrics on total counts and the visits in which each data element is recorded, supporting prioritisation of data rich variables for downstream quality control and analysis.

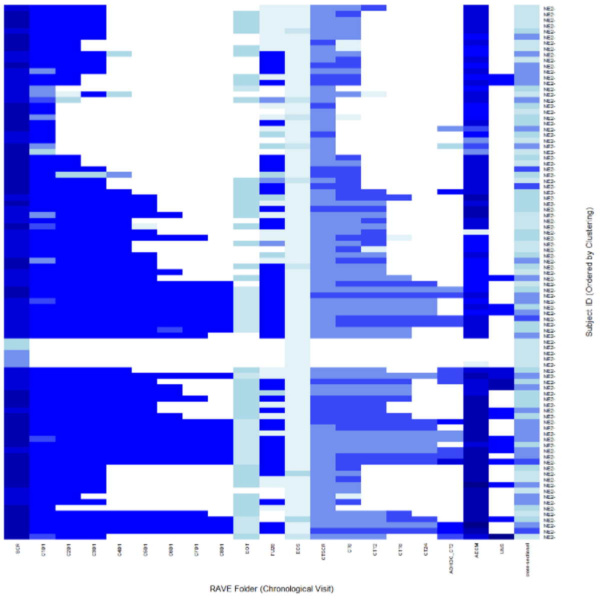


*Supplementary Figure 1 Participant-Visit Non-Missing Field Counts Heatmap.*

*Focused Data Quality Checks (Script 06)*

Performs automated validation of data against a predefined Data Dictionary, focusing on critical data elements. Validation rules include mandatory field checks (identifying missing values), data type enforcement (e.g. numeric, date, categorical), range validation for numeric variables, and coded value verification against predefined permissible values. Additional checks include statistical outlier detection and identification of infrequent or anomalous categorical entries using frequency thresholds to detect potential data entry errors. The workflow generates a detailed issue-level report capturing all identified discrepancies with contextual metadata (form, variable, record, and issue type), alongside a summary report providing aggregated counts of data quality issues to support targeted review, correction, and audit readiness.

### Phase 3: Data Ingestion

This phase populates the core Structured Query Language (SQL) database tables using a long format key–value architecture, in which each data element is stored as an individual record linked to participant and visit identifiers.

*Core Table Population (Scripts 08-10)*

The initial ingestion establishes the participant and visit records that act as the foreign key backbone for all clinical data. For participant records, Script 08 loads the master Participant-Visit units of analysis and the chronological sequence to populate the core subject table. It maps the anonymous participant ID to a unique anonymised participant identifier entry and assigns the numeric Visit number, performing a final row count check to validate the bulk write operation. For visit records, Script 09 links the chronological visits with a human-readable Description (e.g. “Baseline”, “Follow up 1”). This data is written into the clinicaldata table and includes a deduplication check to ensure that the script can be re-run without creating duplicate visit entries.

Script 10 handles the ingestion of all remaining clinical variables:

- **Variable Selection:** The script relies on the manually maintained COSMOS flag, dictated in the Data Dictionary to filter and select only those variables intended for ingestion into the database.
- **Data Transformation:** The selected variables are retrieved from the clinical data CSVs, merged with the foreign keys from the subject table, and transformed from their wide format (variables as columns) into a long format (each row contains a single variable, value and description).
- **Final Structural Enforcement:** A final deduplication check is performed just prior to the bulk write operation to the clinicaldata table. This process enforces structural integrity by ensuring all records link to a valid participant and visit foreign keys, acting as the final technical firewall against structural failures.

### Phase 4: Post-Ingestion QC Certification

This final phase provides the necessary human and automated audit trails required for data release.

*Manual QC Report (Script 11)*

Implements a random sampling strategy (e.g. *n* variables across *k* participants). The output CSV functions as an audit trail by providing full contextual detail, including anonymized participant identifier, Timepoint, Clinical_Variable and Value, enabling the data management team to cross-reference these sampled data in the live COSMOS database against the source data to certify data integrity prior to data release for discovery.

### COSMOS RShiny Tools

Two R Shiny applications provide direct interaction with data stored in COSMOS for users without SQL or R programming experience:

(i) the **COSMOS Interactive Query Tool**, for retrieval of curated clinical data using predefined SQL templates, and
(ii) the **COSMOS Exploratory Analysis Tool**, for interactive exploratory analysis of linked omics and clinical data.

### Query Tool

A demo instance, example files and installation instructions are all available on the GitHub repository: https://github.com/at2e19/SCTU_COSMOS_DQDV_Shiny/blob/main/vm_query/app.R

*Query retrieval*

The query tool allows for data exploration through no-code free-text search. This is done via an inverted-index which maps user-entered keywords to available SQL templates. Where multiple templates match a query, users can select the most appropriate option from a drop-down list (Supplementary Figure 2).

*Parameter entry and execution*

After query selection, the application dynamically generates the required parameters for execution (for example, StudyID or anonymised participant identifier). Query results are displayed in an interactive table and can be exported as CSV files for downstream analysis or reporting.

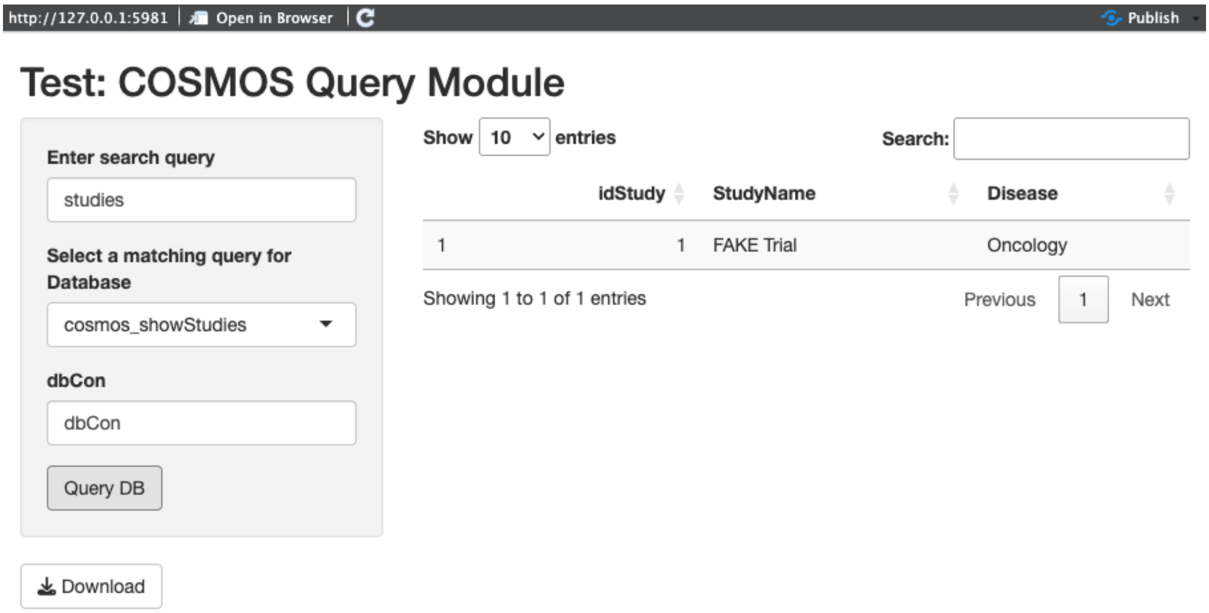


*Supplementary Figure 2: User Interface of the COSMOS Query App. Highlighting the use of free-text mapping to a pre-specified query in the inverted index and the subsequent output from the SQL database.*

**Exploratory Data Analysis Tool**

The application can be found here:
https://github.com/at2e19/SCTU_COSMOS_DQDV_Shiny/blob/main/vm_ml/app.R

The Exploratory Analysis Tool enables exploratory and statistical analysis of omics data linked to curated clinical metadata.

The application uses omics data stored as an RDS file containing an ExpressionSet object. Users specify the relevant StudyID and DatasetID to link the uploaded omics dataset to clinical metadata stored in the SQL database.

Users may select one or more clinical variables to merge with the omics dataset. Selected variables are appended to the phenotype data of the ExpressionSet object and used for downstream stratified visualisation and modelling.

*Implemented workflows*

The application provides the following interactive analysis workflows:

- Summary statistics for omics data (mean, median)
- Principal Component Analysis
- Hierarchical clustering
- Differential expression/abundance analysis
- Random Forest-based feature ranking

*Differential Expression/Abundance analysis*

For differential expression/abundance analysis, the user selects two clinical subgroups for comparison. The application then fits a linear model using limma and returns a results table containing log fold-change, average expression/abundance, raw p-values, and adjusted p-values. Results can also be visualised as a volcano plot (Supplementary Figure 3).

*Feature selection*

The Random Forest module optionally combines omics and clinical variables for supervised feature selection. The resulting variable-importance table and plot (Supplementary Figure 3) provide a ranked summary of predictive features associated with the selected outcome.

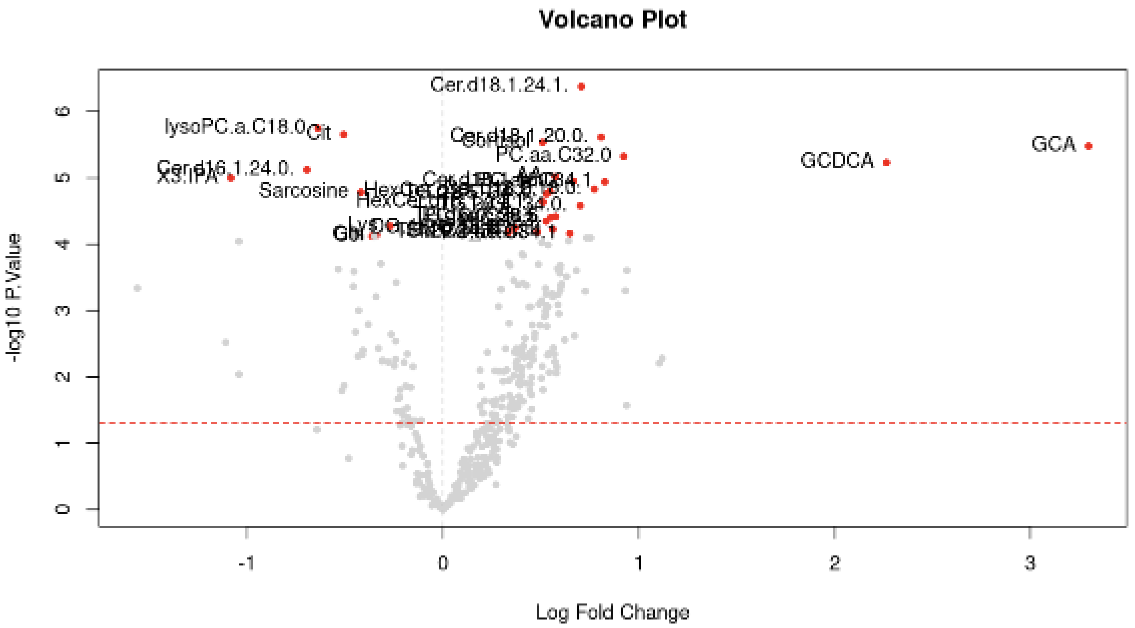


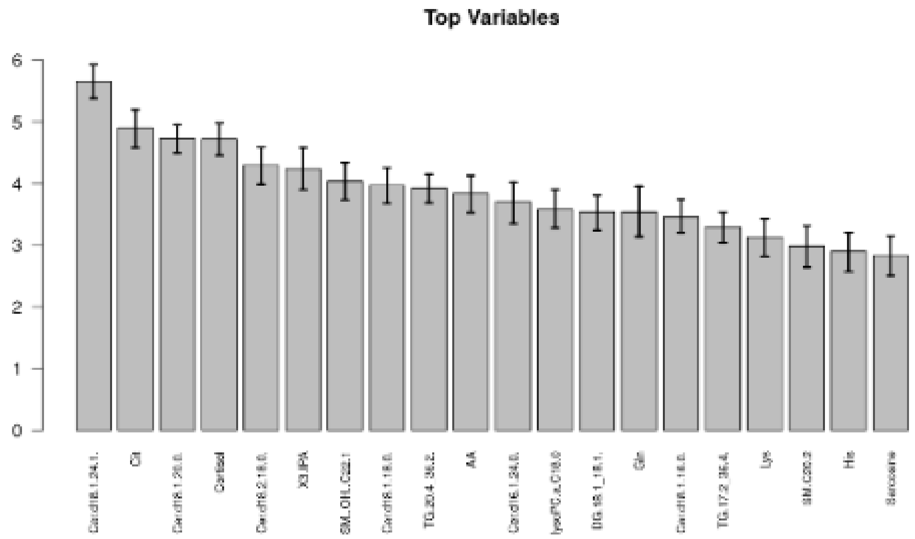


*Supplementary Figure 3: Exploratory Module Output Examples. (Top figure) A volcano plot identifying differentially abundant metabolites. (Bottom figure) A variable-importance plot providing a ranked summary of predictive features associated with the selected outcome from the Random Forest module which enable user to optionally combine omics and clinical variables for supervised feature selection.*

**Infrastructure and Security**

To ensure secure and reproducible access to the data warehouse, the applications are built upon an environment-aware architecture.

*Path Configuration (globals.R)*

The system automatically detects the operating environment (e.g. Windows or Linux) and sets the root directory paths (gcShareRoot) accordingly. This ensures a seamless transition from local testing to production deployment without requiring manual code alterations.

*Secure authentication and database connectivity*

Access to the underlying COSMOS SQL schema is strictly mediated through system-defined Data Source Names (DSN). This architecture ensures "behind-the-glass" database access, safeguarding sensitive clinical metadata while maintaining a persistent and secure connection for data retrieval and joining. All requests to the R shiny applications are required to come from an authenticated and authorized client. Authentication and authorisation are handled by an Entra ID app registration to take advantage of enterprise security features with authorised users represented by an assigned security group.
